# CSynth: A Dynamic Modelling and Visualisation Tool for 3D Chromatin Structure

**DOI:** 10.1101/499806

**Authors:** Stephen Todd, Peter Todd, Simon J. McGowan, James R. Hughes, Yasutaka Kakui, Frederic Fol Leymarie, William Latham, Stephen Taylor

## Abstract

The 3D structure of chromatin in the nucleus is important for gene expression and regulation. Chromosomal conformation capture techniques, such as Hi-C, generate large amounts of data showing interaction points on the genome but these are hard to interpret using standard tools. We have developed CSynth, a high performance 3D genome browser and real time chromatin restraint-based modeller to visualise dynamic and interactive models of chromatin capture data. CSynth does its calculations in the GPU hence is much faster than existing modelling software to infer and visualise the chromatin structure which also allow real-time interaction with the modelling parameters. It also allows straightforward comparison of interaction data and the results of third party 3D modelling outputs. In addition we include an option to view and manipulate these complicated structures using Virtual Reality (VR) allowing scientists to immerse themselves in the models for further understanding. This VR component has also proven to be a valuable teaching and public engagement tool. CSynth is web based and available to use at http://csynth.org.

## Background

With the increasing amount of genomic contact-C-based data becoming available, such as 3C, 4C, 5C, Hi-C, Capture C and now Tri-C^1^, there is a need to understand chromatin structure beyond visualizing data on a 2D genome browser or using heatmaps.

Furthermore, GWAS studies^2^ show non-genic mutations to have a role in a number of diseases, including diabetes^3^ and cancer^4^ and to understand the aberrant gene activity caused by changes in genome structure and the resulting enhancer/promoter rearrangements. The advent of sophisticated microscope imaging of chromatin to observe these structures using super resolution microscopy^5^ and electron microscopy^6^ offers the ultimate means of understanding 3D genome architecture but these methods are slow and expensive. Modelling and visualising the structure of chromatin is useful to understand how genome structure in the nucleus affects gene expression and ultimately why phenotypic characteristics occur in different tissues and cell types.

## Results

The structure of chromatin in a cell is complex, dynamic and not well understood. Methodologies to model such a structure using Hi-C use advanced physics simulations that take several days and large compute resources ^7,8^. An alternative is restraint based modelling^9–16^, which uses Interaction Frequency (IF) to generate spatial restraints and tries to find the optimum model satisfying these restraints based on an objective function^17^. CSynth is unique in that the user can upload such predetermined models of 3D coordinates (which we call ‘xyz input data’), but also allows on the fly visualization of interaction data using restraints in a standard web browser (Chrome, Firefox, Safari or Edge) using the computer’s GPU. Simultaneously, CSynth shows a heatmap view underneath the model allowing visualisation and understanding of Hi-C interactions and their relationship between 2D to 3D space. For example, using CSynth modelling Topologically Associated Domains (TADs) can be quickly identified including other interaction outliers. In addition, we expose various modelling parameters that allow the user to adjust forces on the model and directly see the results. This interactivity is unique to CSynth and allows a much better understanding of the effects of the parameters. The easy creation of such views allows different hypotheses to be tested and then compared to more computationally expensive polymer models or 3D images if these are available.

In Figure 1 we show data generated from Tri-C data set ^18^ at the alpha globin region in mouse captured in erythoid cells. Clearly visible is the chromatin looping of the α-globin (mm9, chr11:32,000,000–32,300,000) self-interacting domain (SID). The coloured sections of the model represent genes loaded as BED format and the ChIP-Seq data uploaded as WIG format. Later sections discuss how CSynth can compare the IF data with the modelled conformation, both visually on the heatmap and by statistical comparison.

**Figure 1.**
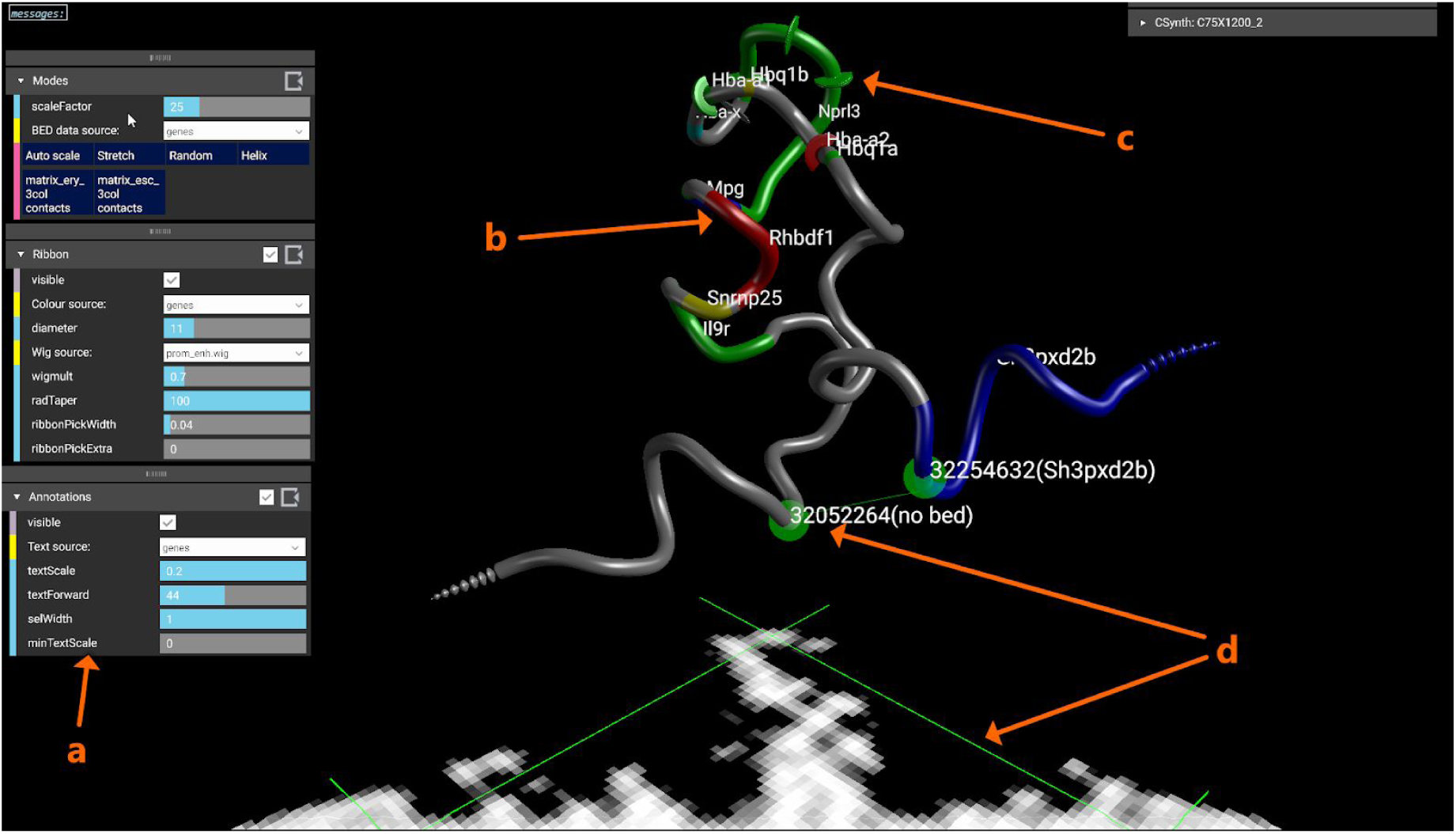
Overview of CSynth features. CSynth allows both the heatmap and model generated conformation capture to be loaded and inspected. “a”: Shows a sub selection of menus that can be used to adjust the visualisation. “b”: Shows genes uploaded as *BED* (Browser Extensible Data) format file for the region. “c”: Visualisation of H3k4me1 data that has been uploaded in *WIG* (wiggle) format, the size of which may be adjusted using the *wigmult* parameter. “d”: The pair of green lines represents a point selected on the heatmap and the corresponding point on the 3D model which is useful to investigate patterns leading to structures seen in the heatmap.

In Figure 2 we show an example of loading a large Hi-C data set at 2 kb resolution from *Schizosaccharomyces pombe* Chromosome I comparing the difference between mitosis and interphase states^19^ using the dynamic GPU modelling built into CSynth. To find the parameters for modelling, we used three dimensional distance between certain chromosomal locations^20^. In interphase (Figure 2a), chromatin fibre forms a characteristic structure and its telomeres, the ends of the chromatin fibre, locate in the vicinity as expected from Rabl orientation within the interphase nucleus in *S*. *pombe* ^21^. Additionally, it has several interesting folding patterns and looping that are not obvious in the heatmap view. In mitosis (Figure 2b) one can see the structure is more compact folding into the characteristic structure and each arm becomes individualised. Clicking on an area of the 3D model shows the chromosome position on the heatmap. Similarly clicking on a peak in the heatmap which would correspond to a frequent interaction of chromatin, and shows the corresponding region in the 3D model. This allows the user to have a direct understanding of the relationship of the heatmap with the 3D model. The fact the structure curved around towards itself at the ends was unexpected. Consideration of this showed that they came from contacts at very long backbone distances; eg contacts between the two ends. Looking deeper showed that there was significant noise in the contact data for high backbone distances which led to greater insight into the source data and better understanding of artefacts. These effects can be bypassed using the parameter MaxBackboneDistance to ignore noisy items.

**Figure 2:**
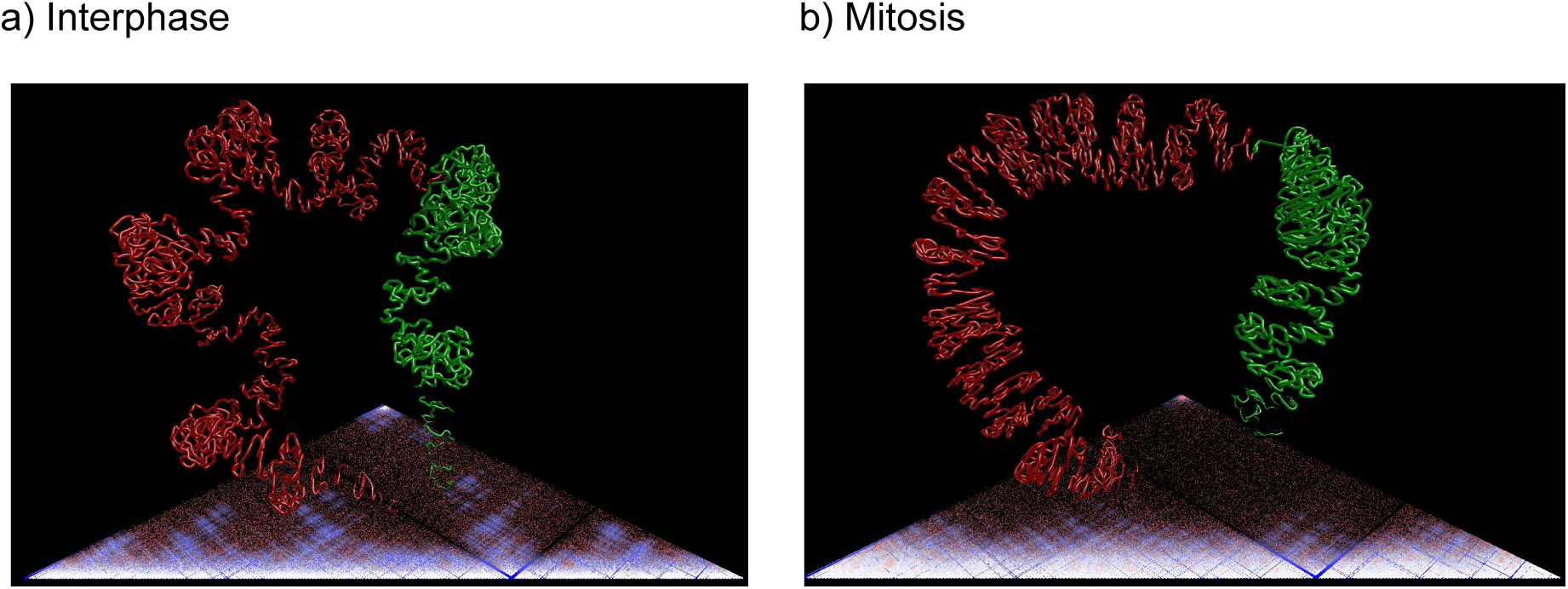
*S. pombe* chromosome I comparing difference between mitosis and interphase states. Red and green colouring show the two arms of the chromosome.

## Virtual Reality

Virtual Reality (VR) offers an improved way of viewing and interacting with these complex datasets allowing new perspectives on the data that would not be afforded via a 2D screen. For example, complex chromatin loops can be observed at different points of view when comparing the erythroid (ery) versus embryonic stem cell (esc) at the alpha globin locus. The orientation of the model is controlled by the mouse or via a touch screen (depending on what is available). The VR mode is implemented using WebVR and available in Chrome and Firefox browsers activated by pressing the F2 function key. The experience is tailored to use with the HTC Vive headset which comes with two (hand) controllers. The first controller also doubles as a way to rotate the model in 3D space, which is much more intuitive and flexible than using a mouse. A slider on the dashboard allows the user to zoom into the model and go inside the chromatin structure. The menus are available in VR and may be customised according to the amount of functionality required. We find that VR is extremely valuable in public engagement activities and teaching, and provides good flexibility for setting up demonstrations.

## Availability

CSynth can run directly in a web modern browser on all operating systems (including tablets). Data can either be uploaded to the server for later use and for sharing, or can be directly drag-dropped from the local file system for quick viewing.

On laptops with both integrated and discrete graphics it is worth making sure that the discrete graphics are used. There is no significant performance difference between browsers once a session is running.

## Visualisation

### Views

The main views provided by CSynth are the ribbon and heatmap views previously mentioned. It also has a view where each particle is seen as a sphere. This can be helpful to see how the dynamics is placing the particles along the 3D ribbon. A “history trace” view shows a trace of the recent dynamics history of each particle (typically generating a folding surface). This helps visualize the dynamics, and can be used to provide a static image of a dynamic system.

### Colouring

The main views can be coloured by different properties. These may be derived from *BED* (Browser Extensible Data) annotations; colouring according to the colour defined in the file (if present) or a standard sequence of colours (if none specified in the file). Rainbow colouring colours in a spectrum along the ribbon brings out the intricacies of its folding such that areas that are geometrically close but distant in the genome can be identified by their contrasting hue.

Points on the heatmap are derived from two particles. We can colour according to the current distance between them, the distance in any xyz input file, or the IF from any input matrix file (Figure 3). We can also colour according to a combination of the single particle properties. For example, we can tint the heatmap by the bed colouring used on the ribbon; this gives immediate visual correlation between the ribbon and heatmap.

**Figure 3:**
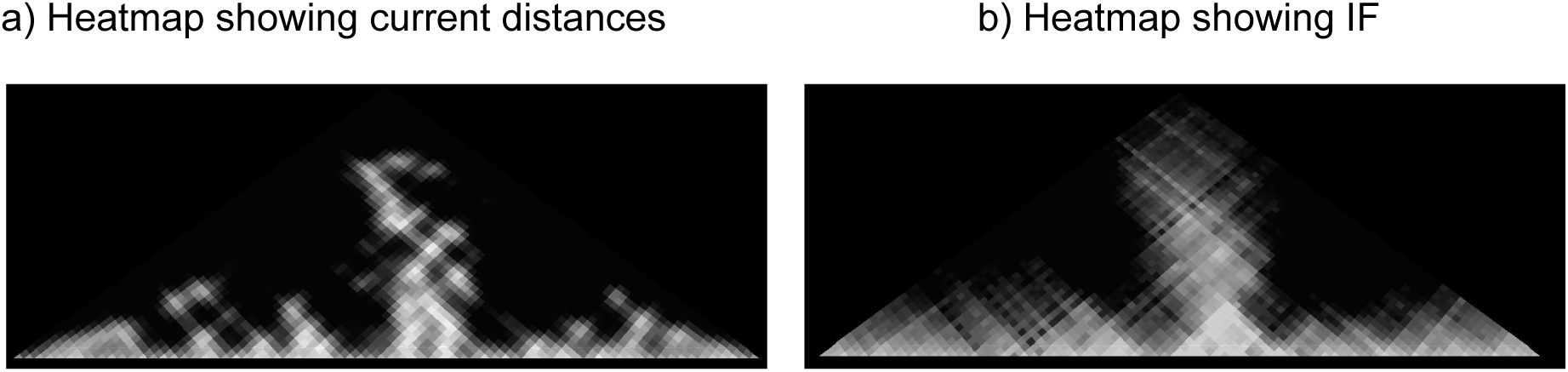
heatmap showing different properties of the matrix file.

**Figure 4:**
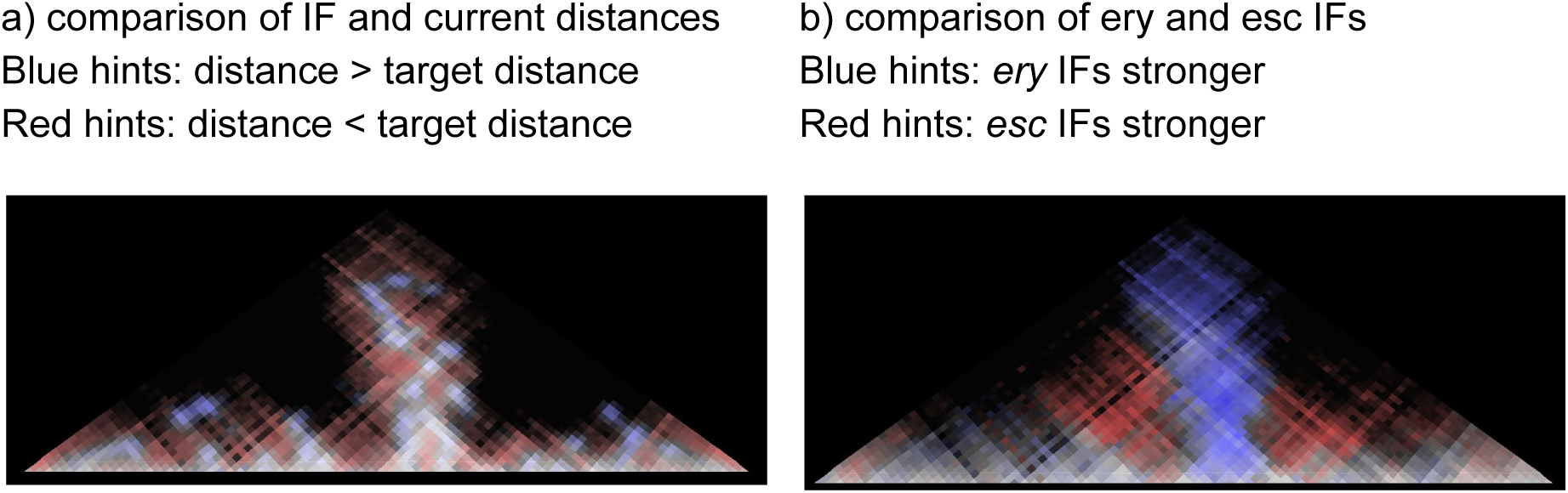
Heatmap showing comparison of different properties.

Colouring may also be based on comparison of properties (see Figure 5). For example, a comparison of IF and current distance on the heatmap shows how closely the current conformation agrees with the IF. Matching areas are shown on a grey spectrum, from black for particles currently distant and with no expected contact, to white for close particles with high IF. Non matching areas are coloured; red shows regions that are currently close despite having low IF; blue for regions distant in the current conformation despite high IF.

**Figure 5:**
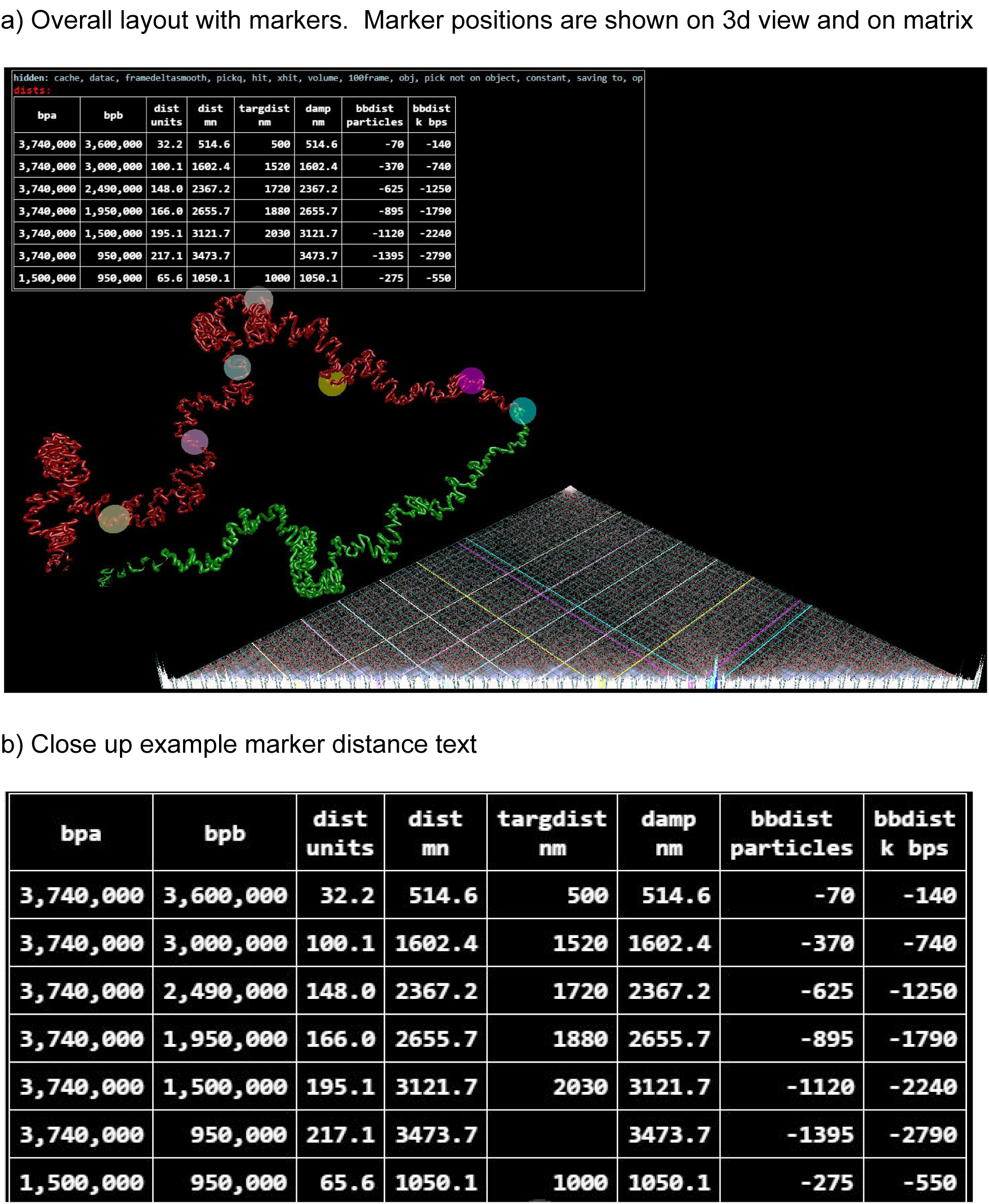
Markers and distances provide waypoints that can be used to compare model distances with real world measurements.

IF and distance cannot be compared directly. We convert IF values to a target distance (more details in the following Modelling section), and then map to a 0 to 1 scale (1 for close or strong).

Similarly, colouring can be used to compare two contact maps, current distances versus xyz input derived distances, and so on. This is used to see the differences between the contact maps for different states; and for visual comparison of different models, for example where we have loaded different xyz files from external modellers.

### Markers

The user can place markers on specific positions in the model, and CSynth will monitor and display distances between them. It can also show known experimental distances, such as from 3D-FISH experiments, so that these can be compared against distances with the current model. The markers are highlighted on the 3D model and the matrix (see Figure 5).

## Further useful visualizations and experiments

CSynth has many ways of showing features that are not naturally occurring, but which help the user understand the data and its implications.

We can transition between the conformations derived for various input files. We switch the drivers of the dynamics from one set of input data to another, and the dynamics performs a smooth transition between conformations. Such transitions are not part of real biology, but seeing the transition process highlights the difference between the conformations in a way a comparison of two static conformations cannot and that is more meaningful than simply interpolating geometric positions. These transitions may be between the same model inputs (whether raw / normalised IF, or derived xyz inputs from external modelling systems) to understand the differences between different states. In the example shown in figure 1, we show interphase and mitosis states, but other examples could be models from different tissues or cell types as shown in figure 2. Also we may see the transitions between the same state for different modelling systems to get a feel for how the models agree and how they disagree We can also load raw and normalized matrices to compare before and after normalisation and its effect on the structure.

When one or more data sets are loaded, CSynth presents buttons associated with that data: a “positions” button sets the exact xyz values from the file, and a “distances” button sets up dynamics based on the distances. The dynamics model means that transitioning between two sets of distances naturally tends to keep a good consistent orientation between them, even where the positions themselves are not registered. The forces within the dynamics give a fairly natural transition, not just a straight-line interpolation. In general distance based models cannot simultaneously satisfy all distance constraints; where the distances are derived from positions the model can establish the conformation that matches all distances.

The model can be set to some initial ‘wide’ view and allowed to fold. As the folding happens, the TADs form before the complete packing occurs. The TAD can be difficult to see in a fully packed model even when highlighted, thus the interactive folding gives a better concept of TADs, as well as the final static packing giving some idea of real configuration.

We can exclude most particles and gradually reveal them, incrementally spinning out the strand with the dynamics forcing folding as the full model is revealed (see Figure 6a)

**Figure 6:**
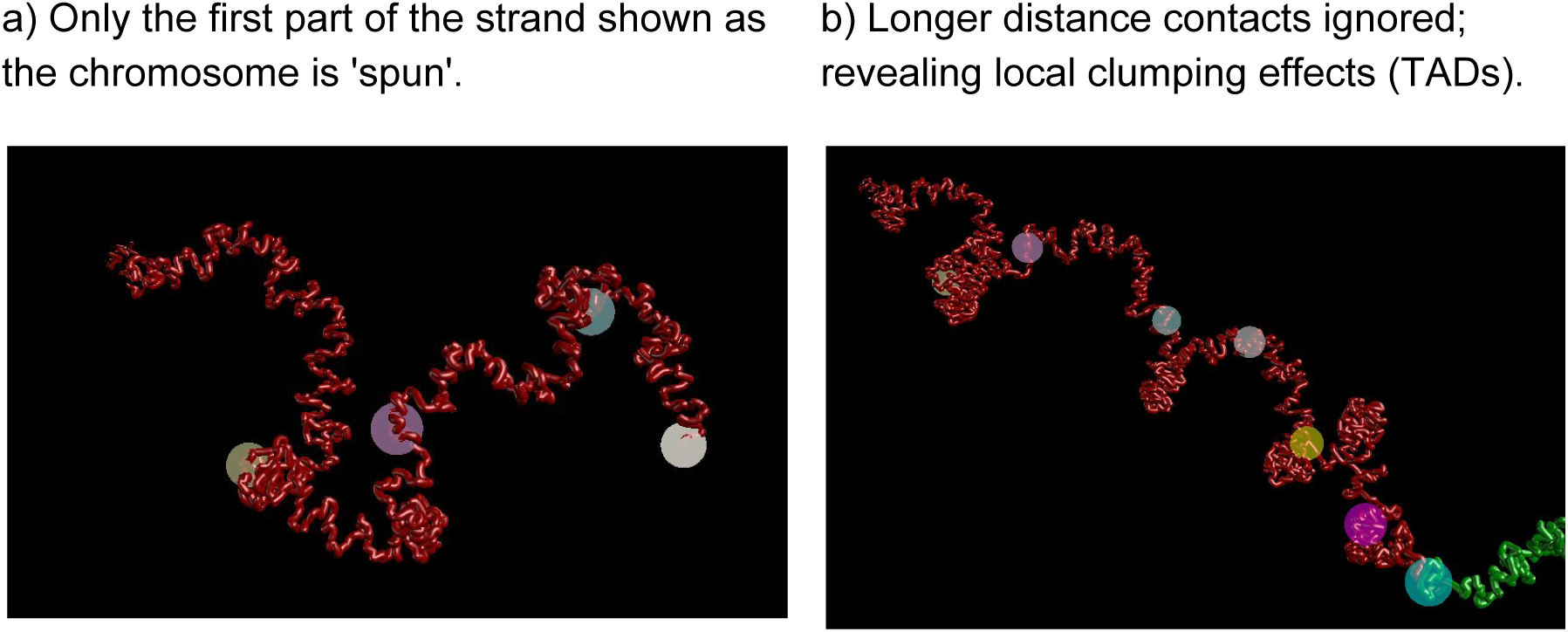
Special transition effects.

Alternatively, we can limit almost all inputs except for those over very short backbone distances, and then gradually enable them over long distances. This helps the user separate out the finer detail derived from short backbone distance information, and the broader scale conformation that results from longer backbone distance contacts (see Figure 7b). The effect is similar to the dynamic folding but with more direct user control, and easier for diagram production.

**Figure 7:**
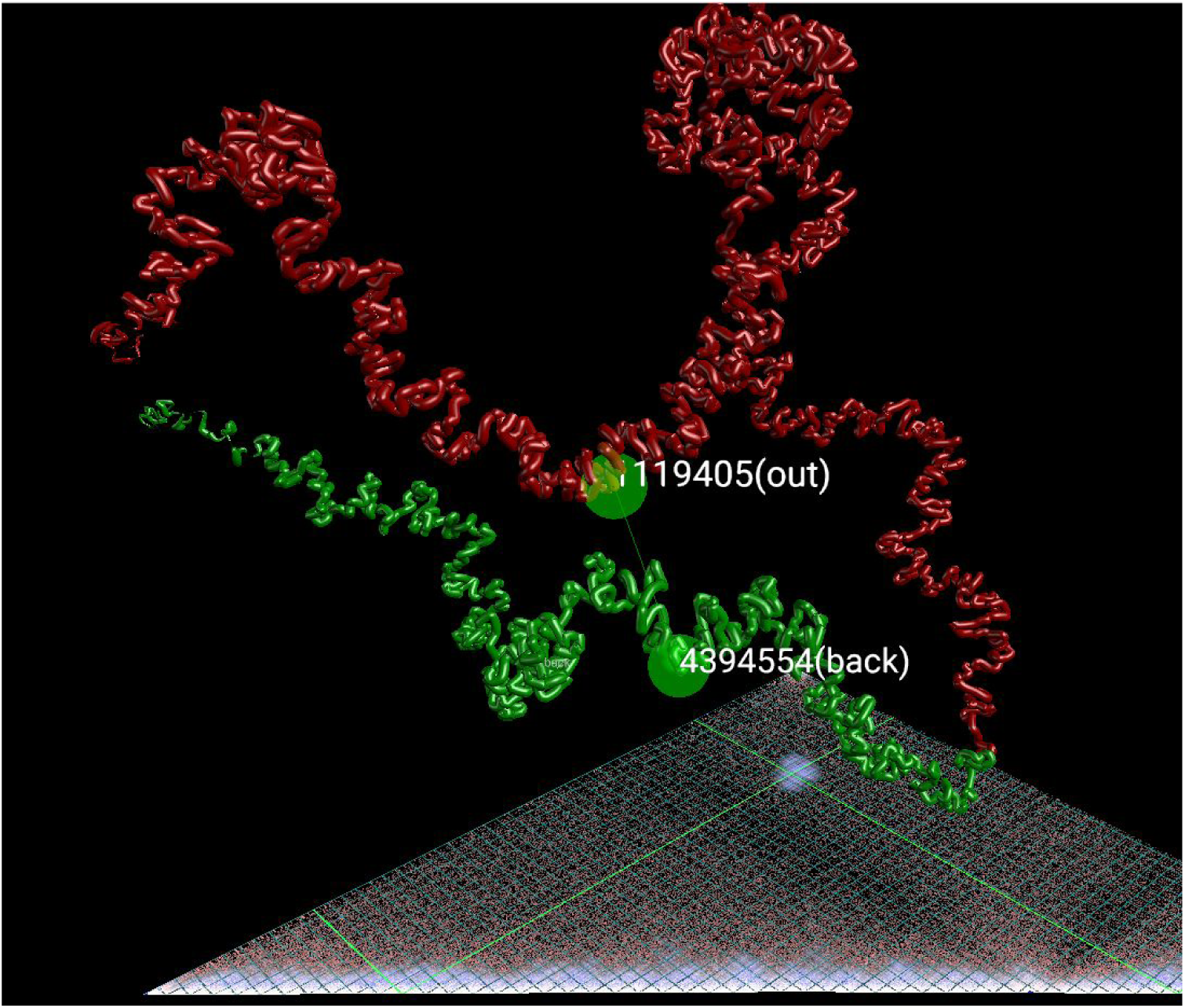
Boosting. A point on the heatmap is selected; it is highlighted in green on the matrix and the 3D view. Contacts in the nearby region are boosted so the dynamics brings those regions closer. The white region on the heatmap indicates the stronger contacts and closer distance.

The ability to ignore longer distance values can help handle some noise in the data. For example, the yeast modelling always bent the mitosis model almost into a torus. This conflicted with other data not available to our model. We realized that the main cause was false/noise IFs recorded in the data between the start and end of the chain due to mapping issues in the Hi-C data within repetitive regions. Filtering out these IFs within the interface permitted a more expected view. The cutoff can be changed interactively with immediate effect on the dynamics and quick effect on the configuration. This interactivity gave the scientist more insight into the implications of the data.

We can give viewers a clearer orientation on the object by attracting the two end particles to points on the x axis in what we call a ‘skewer’ effect. For example, when we model a locus we can simulate a particular pull upstream and downstream of that locus because physical forces such as occlusion with other regions of packed chromatin in the nucleus do not allow the two ends of the region to come together. This is often necessary in the dynamic modelling to prevent the two ends of the region of interest coming together by the sheer number of interactions in that region.

In an extreme form of this we can completely pull the attraction points apart, freeing non-backbone inputs to completely unwrap the folding. This has the effect of straightening the entire region of interest and shows how long the chain is (even though it only unfolds to particle level, not to base pair level) and looks more like a traditional 2D genome browser view.

It is clear from the IF data that no single configuration accurately represents what is going on; only many different configurations can ensure that all observed IFs are possible. We can explore a range of configurations by vastly strengthening the IFs between two small regions of the strand; this ensures the associated particles actually do interact. More details are included in the Dynamics section below. Moving the enhancer region, for example by moving the mouse over the heatmap view, gives a much clearer idea of the configuration variance (see Figure 7).

## Dynamics and modelling

Rather than simulating sophisticated molecular dynamics CSynth uses simple forces to seek conformations that best satisfy the known IFs. This builds on the work we used in FoldSynth^22^ which is software we developed for interacting with protein structures. For example, unsatisfied IFs can have an effect on very long distances. This allows simpler (to compute) dynamics. Our dynamics are inspired by previous work by Jefferys et al. (aka Poing^23^ largely based on spring-like dynamics). Some of the forces used in our dynamics may be related to real physical forces but the relationship is usually indirect; our dynamics are better thought of as an emulation rather than a simulation or modelling. The various forces we have built in CSynth are detailed later in this section.

Our dynamics work directly from IF or distance map inputs, which are held in sampler buffers on the GPU. The modelling system is based on particles, which are represented using the size of the fragment from the capture experiment. The particles generally match the Hi-C bands one to one, but we permit the use of multiple particles per cell for more refined modelling. The particles are assumed to be joined in a backbone chain (or chains). The modelling operates in conventional Newtonian dynamics steps; in each step an overall force is computed on each particle; the force is applied to the velocity, and the velocity used to compute a new position.

The number of particles is limited by GPU texture constraint which is typically, as of writing, 16000 particles. In tests, we have resolved models of 6284 particles (3 chromosomes of yeast at 2k resolution) in a few seconds on an NVidia GTX 1080, and 2258 particles (yeast chrII at 2k resolution) on a laptop with integrated Intel 4600 GPU.

When modelling large chromatin regions we avoid using Hi-C data that is from centromeric or telomeric region since the sequence data generally maps incorrectly in these regions and do not reflect real interactions.

### Pairwise forces

We have several pairwise forces. The essence of the emulation is the balance between these forces. All these forces depend on current distance between the particles (**len**) and act along the axis joining the particles.

#### xyz input based forces

This is the main force for handling xyz input, which may be in the form of white space separated .xyz files, or .pdb or .vdb files. Strictly this a distance force, which when applied attempts to bring the particle pairs to a given length apart. The distances are computed from input file positions (**targlen**).

~~~
pseudocode 1
     force = (len - targlen) * xyzforce / targlen;
~~~

**xyzforce** scales the force. Division by **targlen** weakens the effect of long distance pairs.

Additionally, we can ignore all pairs where targlen is bigger than a value **xyzMaxDist.** We use this more in protein modelling to visualize protein folding based only on close contacts.

The xyz forces force conformation according to the input data where all other forces are turned off. The other forces can still be used to distort the original data for visualization or what-if experiments.

#### Contact input based forces

The main contact force is an attraction force based on the IF from the input data (**contact**) and scaled by **contactforce**.

~~~
pseudocode 2
     force = contactforce * contact * len
~~~

This works to balance the following global pushapart force so that particle pairs with stronger IFs are brought closer than pairs with weaker IFs.

Input contact data may have simple noise reduction applied by subtraction of a threshold **contactthreshold**, with negative results set to zero. We can also filter contacts for pairs with a backbone distance (relative to total backbone length) greater than threshold **maxBackboneDist.**

Hi-C data is often at a relatively low resolution, so dynamics based on each Hi-C region being a separate particle enforces rather limited conformations. By modelling with more particles than Hi-C regions we get an idea of possible conformation within the region, and how regions interact. An **expand** factor defines how many dynamics particles will model each Hi-C region.

#### Global pushapart force

The global pushapart force works to make all particles repel each other.

~~~
pseudocode 3
     force = -pushapartforce * pow(len/powBaseDist, pushapartpow);
~~~

**pushapartforce** is the basic scaling factor. The pushapart force is attenuated according to distance by power **pushapartpow** (usually negative), Figure 8.

**Figure 8:**
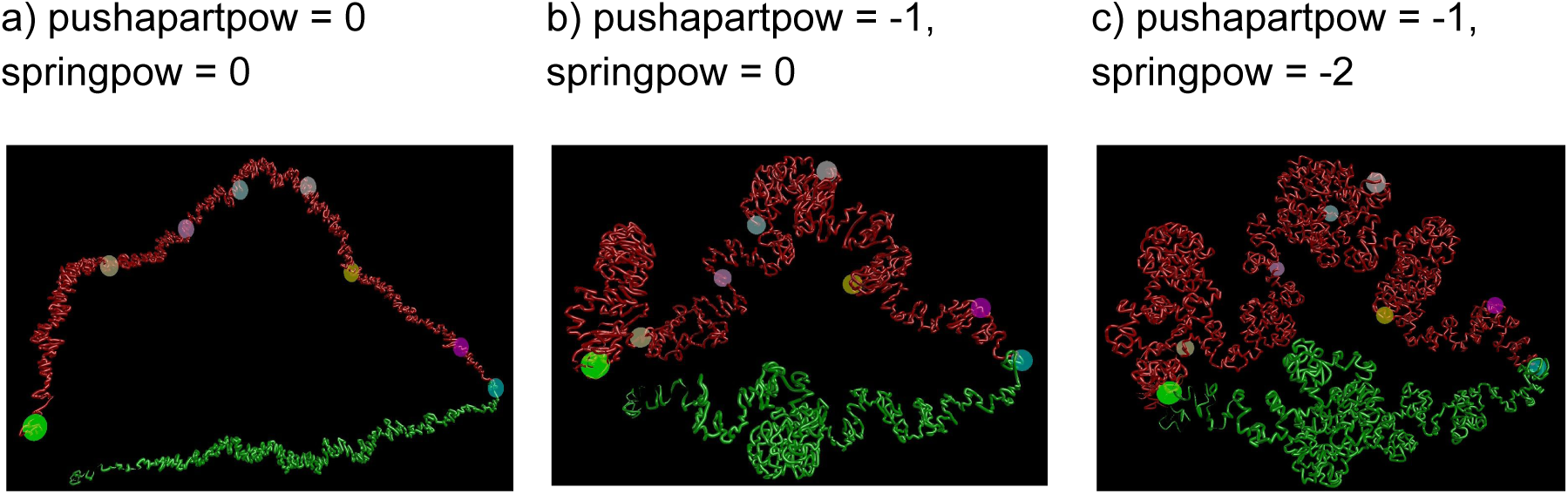
Showing the effects of changing modelling parameters. As forces fall off over distance with negative values this allow more clumping to occur,

The main purpose of this force is to balance the IFs to produce effective contact-based conformations. Boosts to the pushapart (available with ley ‘P’) can also break the dynamics out of local minima.

**powBaseDist** affects the scaling of this attenuation so that the forces at this distance remain constant as the power itself is changed. It is typically set to the approximate overall size of the conformation. Changing the power therefore changes the contact/pushapart balance and thus the detailed conformation of the object, but does not significantly change its overall size.

#### Local pushapart force

A local ‘pushapart’ force prevents particles getting too close; this is very strong when particles get too close and falls in an S-curve to 0 over a short distance (**pushapartuse**). **pushapartuse** applies separately to particles close along the backbone controlled by **nonBBLen**; this can be thought of making the particles ellipsoids rather than spheroids.

~~~
pseudocode 4
     force = -pushapartlocalforce * (1-smoothstep(0.5, 1, rlen))
     *where:*
     bbd: backbone distance between the particles
     nonBBLen: max bbd to consider as related by backbone
     pushapartuse = min(bbd, nonBBLen) : ‘target’ minimum length
     rlen = len / pushapartuse: ratio of current len to target
~~~

#### Backbone force

The backbone force adds an extra xyz style force to bring particles along the backbone to within a unit distance.

~~~
pseudocode 5
     force = (len - 1) * backboneforce;
~~~

This reduces some distortions when transitioning between different states. It can make the conformation appear smoother and more regular (more of an aesthetic than scientific value).

#### Lorentzian force

This applies a force computed as a derivative of the Lorentzian cost function^17^. A special case replaces contact for backbone pairs with the maximum contact value.

~~~
pseudocode 6
     float d = m_k * pow(contact, -m_alpha); // target distance
     float dd = d - len;
     float dem = m_c * m_c + dd*dd;
     gforce += m_force * contact * −2 * m_c * m_c * dd /
    (dem*dem);
~~~

### Single particle forces

Some forces act on single particles. Particles can have forces to fixed points, or be constrained to a fixed point. That can be used to enforce a particular orientation and prevent drift. It can give a ‘skewer’ effect where ends of a segment are pulled out which can help readability and could be potentially used in future simulations to model attachment regions attaching to the lamina of the nucleus.

A damping factor is applied based on 4th power of the velocity, with an absolute velocity limit to help stability, and further damping as the particle approaches a populated region. This does not affect the steady state but it permits the dynamics to execute faster while remaining stable. Noise based on the Langevin model^24^ applies random changes to the velocity. This is available in CSynth. “Boosts” to the noise (available with key ‘N’, see below) can also break the dynamics out of local minima.

### Extra force controls

We have described above the basic forces. There are various ways to manipulate these to explore different aspects of the data.

#### Force falloff

A falloff power factor **springpow** applies to all the global forces.

~~~
pseudocode 7
     force = pow(len/powBaseDist, springpow);
~~~

This changes the balance between forces acting over short and long distances. It could have been applied separately to the various contributing forces, but having it separate means it can be adjusted without altering the balance between contact and pushapart forces and the target distance implicitly derived from those.

### Boost

The “boost” mode allows certain regions of the contact forces to be boosted. The region is an area of the heatmap, so that it connects two regions on the strand. A multiplier centred on the region falls off in an s-curve to 1 at the edge, and this multiplier applies to the contact force above (Figure 8). The region is interactively changed by hovering over the heatmap, with controls for its radius (number of particles affected), and the level of boosting to be applied.

No single static conformation can explain the huge number of contacts seen in a typical Hi-C experiment. Either different samples take up different conformations, or all conformations are highly dynamic, or both. Using the boost finds candidate conformations that exhibit the selected boosted contacts. Holding the selection gives static conformation, and dynamically moving the sections shows a plausible dynamic.

### Statistics

CSynth allows the calculation and display of various statistics for distance comparison. It supports root mean square error (RMSE), root mean square distances weighted by inverse of maximum of the two distances (WRMSE), Pearson and Spearman. RMSE and WRMSE between the wish distances and current distances can be displayed in realtime; the ‘X’ key shows all statistics for that pair; and the full set between wish distances and distances derived from any loaded xyz/pdb files can be displayed with the ‘Y’ key.

## Comparison with other software

CSynth is unique in that it combines the ability to adjust restraint based modelling parameters and visualise the results in real time which makes it extremely convenient to use. Usually packages focus on the modelling and have a separate viewer for visualisation or generate results in PDB format for viewing in a protein structure viewer. For comparison with other packages, we evaluate the visualisation and restraint based modelling in the two following separate sub-sections.

### Visualisations

There are currently several 3D genome browser implementations suitable for looking at 3D chromatin structure. *Genome3d*^*<u>25</u>*^ is a downloadable C++ application which is limited since it requires a computer running the Windows OS and the (manual) installation of software. *GMOL*^*<u>26</u>*^ does not handle Hi-C data, but more recently the author has release *GenomeFlow*^*<u>27</u>*^ which offers a full Hi-C analysis pipeline which is very useful. However, using Java often requires the user to install the relevant Java version as opposed to using the desktop browser, causing a barrier to entry to anyone who wants to rapidly and easily visualise their analysis (data). *Tadkit*^*8*^ is web based and shows a 3D chromatin view in the context of a 2D browser based on *IGV*^*<u>28</u>*^ but there is no possibility provided to show different states (for example in different tissues). Distinctively, the CSynth platform provides a highly interactive, WebGL based user interface that is powerful enough to view many types of capture based chromatin modelling. CSynth allows comparison between different samples and also visualisation and interaction in VR mode. If the user registers, all data is managed and stored in a web based portal. Once registered, users may upload matrices and annotation formats to store details of various experiments and different parameters selected.

### Restraint Based Modelling

#### Comparison vs other software

*Chromosome3D*^*29*^ and *LorDG*^*17*^ have been extensively benchmarked and hence we compare our modelling with the results of these papers. To aid our comparison we have implemented the Lorentzian function reported in LorDG and compared using the synthetic data sets described at https://missouri.box.com/v/LorDG.

We chose one of their experimental groups (chr20 from chainDres25), and loaded the IF data plus their 10 pdb results files, 5 from LorDG optimization and 5 from square IF optimization (see Figure 10). We visualise their results by clicking on the various ‘positions’ buttons. Alternatively, using the ‘dists’ buttons transitions between the results and brings out similarities and differences, especially when using ***history trace*** view. This helps visualize the differences between their LorDG and square results, and the regions of the chromosome that the results indicate are most variable. We note that the “history trace” is best seen time-based. However, one of its strengths is that it displays motion in a static printable image (with correct colouring/background/emphasis).

**Figure 9:**
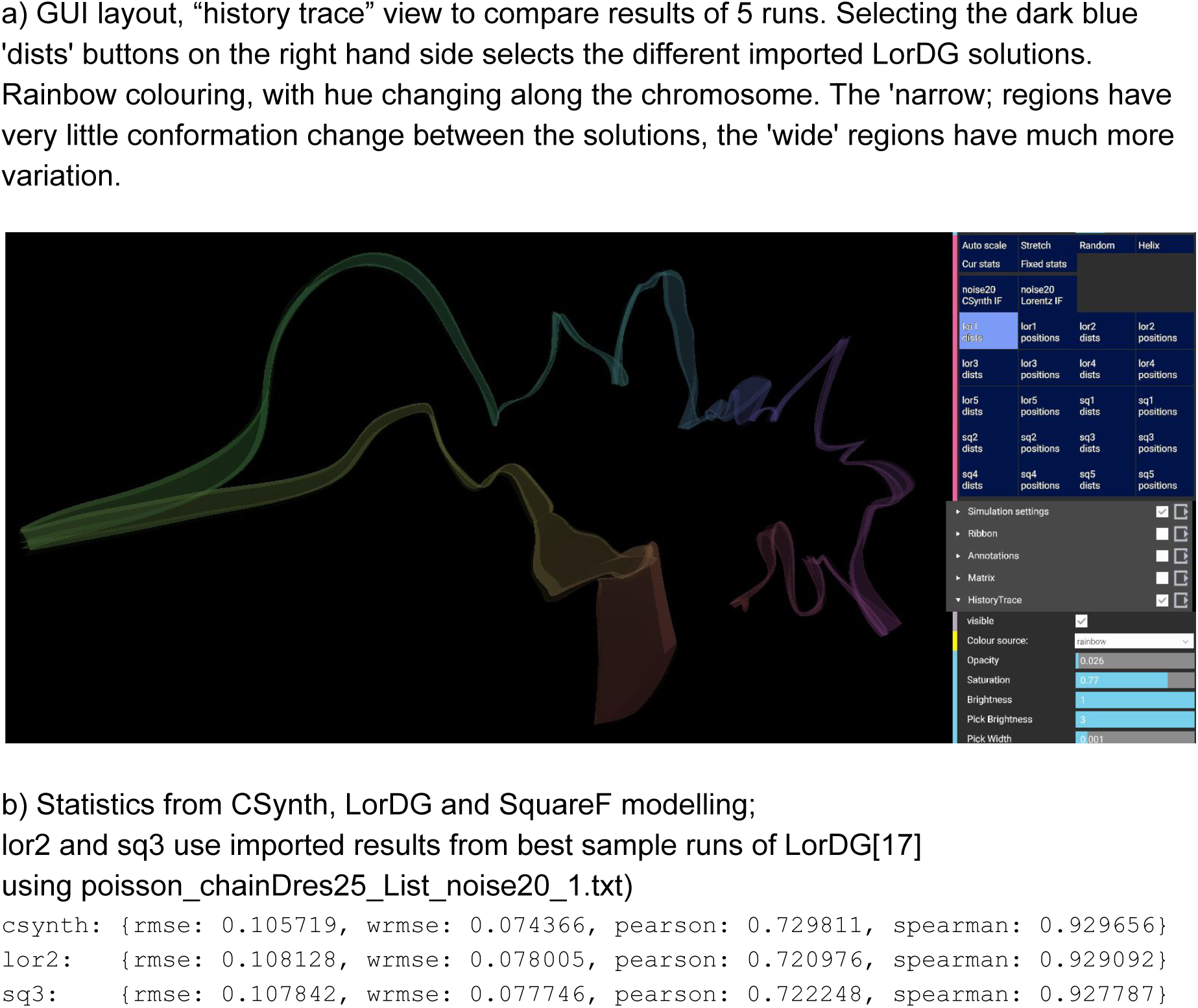
a) Exploration of LorDG data b) comparison of CSynth with other modelling software.

**Figure 10:**
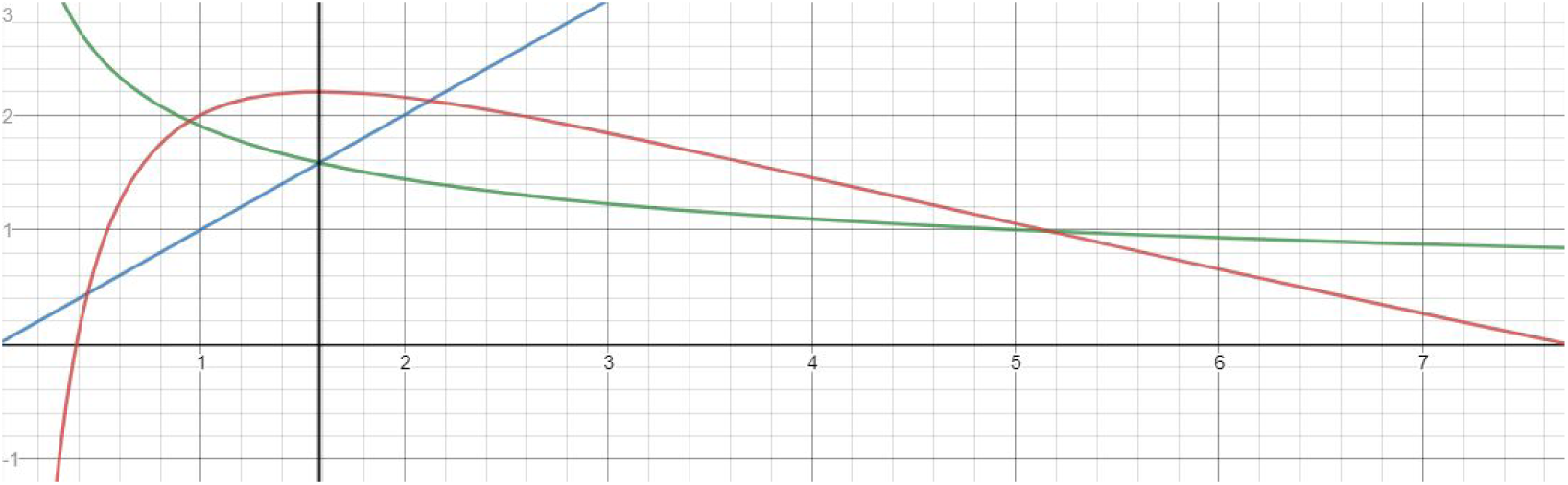
Balance of contact and pushapart forces: force vs distance. Green: pushapart force, blue: contact force, black: implicit target distance, red: equivalent cost function (also allowing for springpow) https://www.desmos.com/calculator/a6rnymfebx

We can also run our own model using the IF, and transition between our models and LorDG result models. Finally we have added forces to CSynth that are very close to the LorDG model and can execute in real time for experimentation with the parameters (such as c and alpha). One can test these forces repetitively using the Stretch, Random or Helix buttons. Running stats on this single experiment indicates that CSynth modelling gives marginally better results than either LorDG or square in Chromosome3D (Figure 9.b).

For almost all modelling systems CSynth provides the ability to load and visualize results and compare them with CSynth models. We intend to permit add-on extensions to run other simple modelling systems within the CSynth framework in real time; this is not feasible for more complex *ab initio* modelling.

### Dynamics and optimization

Though CSynth dynamics were conceived and implemented in terms of forces, the effect is very close to modelling systems that use relaxation or hill climbing to optimize cost functions based on distances between particle pairs. Integrating the effect of our forces gives an equivalent cost function, conversely hill climbing generally differentiates the cost function to provide a hill ascent function. Several other systems use an inverse alpha power falloff for converting IFs to target distances, -alpha is directly comparable to 1 / (pushapartpow-1); see Figure 10.

### Comparison vs hi resolution single cell super resolution imaging

As an example of efficacy of CSynth’s modelling we use the mouse alpha globin locus (figure 11). We load matrices generated from TriC data data^18^ spanning mouse (mm9) at 4kb resolution. Comparing the resulting model with super resolution microscopy generated using RASER-FISH^30^ the chromatin loop or self interacting domain (SID) is clearly visible in the matrix and the model.

**Figure 11:**
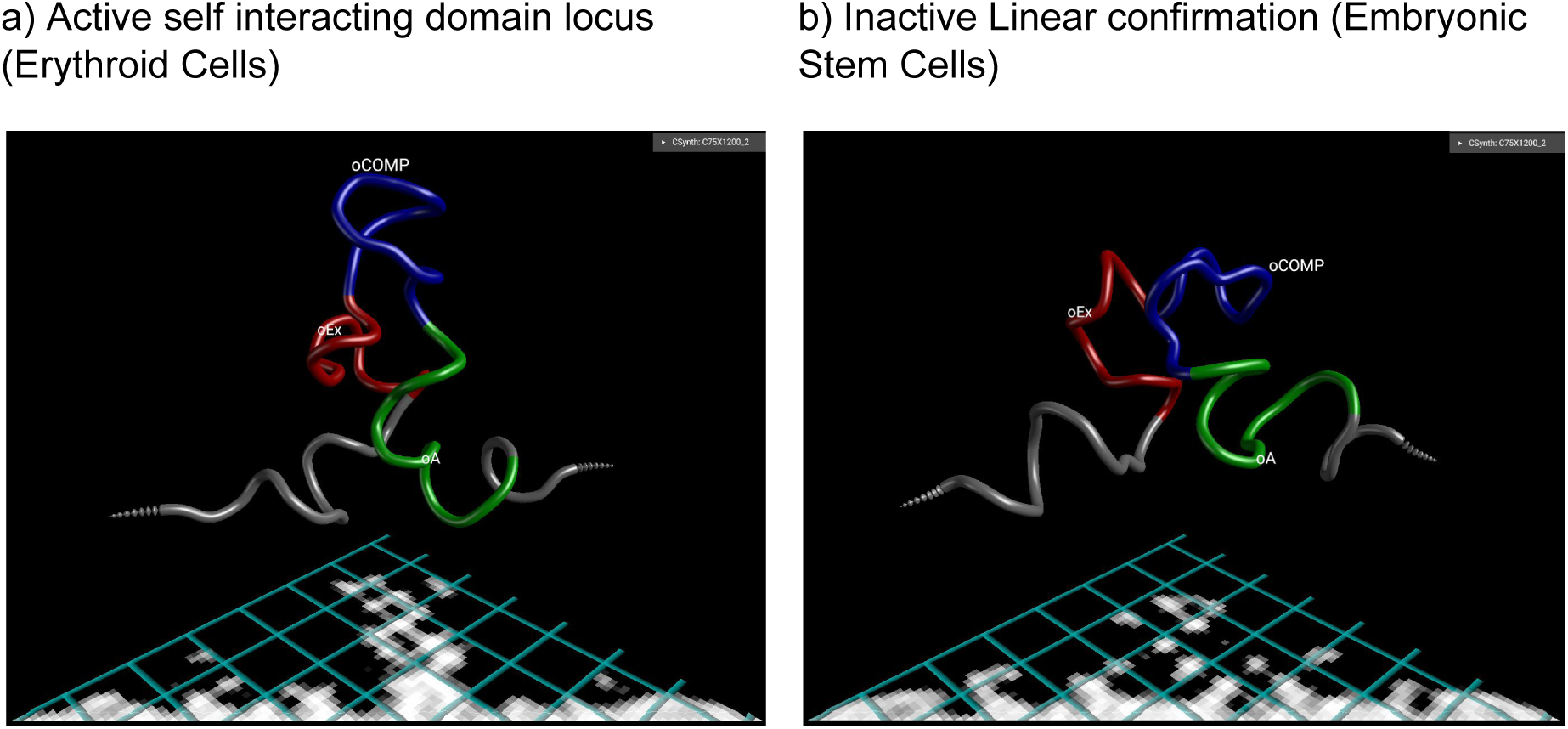
Comparison of the alpha-globin locus in different cell types. The alpha-globin SID (blue region, labelled oComp) when alpha-globin region is active (a) and not extruded when inactive (b).

## Conclusion

CSynth provides an interactive, user friendly and powerful way of visualising chromatin interaction data, by combining model, heatmap and genome annotations in one display in a standard browser. These features are useful when trying to understand the structures of Hi-C and related data. A key improvement in CSynth is that modelling is done on the GPU dynamically. This allows the user to load chromosome capture matrices quickly and vary model parameter values for a better understanding of their effect on the modelling process. Another unique feature of CSynth is the facility to compare models between any number of different samples (e.g. tissues or cell types) or even other modelling systems. Finally, we use VR to view and interact with these complex 3D structures which helps get a better intuition for the 3D modelling and is also useful for teaching and public engagement. We foresee that CSynth will be invaluable to understand the structure and dynamics of more complex data generated from different samples from existing and new C based techniques such as single cell Hi-C^31^.

